# Leveraging Deep Learning to Simulate Coronavirus Spike proteins has the potential to predict future Zoonotic sequences

**DOI:** 10.1101/2020.04.20.046920

**Authors:** Lisa C Crossman

## Abstract

**Motivation:** Coronaviridae are a family of positive-sense RNA viruses capable of infecting humans and animals. These viruses usually cause a mild to moderate upper respiratory tract infection, however, they can also cause more severe symptoms, gastrointestinal and central nervous system diseases. These viruses are capable of flexibly adapting to new environments, hence health threats from coronavirus are constant and long-term. Immunogenic spike proteins are glycoproteins found on the surface of Coronaviridae particles that mediate entry to host cells. The aim of this study was to train deep learning neural networks to produce simulated spike protein sequences, which may be able to aid in knowledge and/or vaccine design by creating alternative possible spike sequences that could arise from zoonotic sources in future.

**Results:** Here we have trained deep learning recurrent neural networks (RNN) to provide computer-simulated coronavirus spike protein sequences in the style of previously known sequences and examine their characteristics. Training used a dataset of alpha, beta, gamma and delta coronavirus spike sequences. In a test set of 100 simulated sequences, all 100 had most significant BLAST matches to Spike proteins in searches against NCBI non-redundant dataset (NR) and also possessed concomitant Pfam domain matches.

**Conclusions:** Simulated sequences from the neural network may be able to guide us in future with prospective targets for vaccine discovery in advance of a potential novel zoonosis. We may effectively be able to fast-forward through evolution using neural networks to investigate sequences that could arise.

## Introduction

### 1.1 Coronaviridae

Coronaviridae are a family of large, enveloped single-stranded positive-sense RNA viruses encompassing alpha, beta, gamma and delta coronavirus divisions^1^ as well as unclassified divisions in the sequence databases. The genome is packed inside a helical capsid and is further surrounded by an envelope. The spike protein forms large protrusions from the virus surface, giving the coronaviruses the appearance of wearing a ‘crown’ under electron microscopy. Coronaviruses have the ability to infect a wide range of different animals and usually cause mild to moderate upper-respiratory tract illnesses, however they can also cause severe respiratory infections as well as gastrointestinal and central nervous system diseases. Coronaviruses circulate among humans and animals such as bats, pigs, camels, and cats^1^. Recent zoonoses include SARS Coronavirus (SARS-CoV), which emerged in November 2002 and became effectively extinct by 2004^2–5^. Middle East Respiratory Syndrome (MERS-CoV) was believed to be transmitted from an animal reservoir in camels in 2012^6^. In veterinary terms, CoV such as porcine epidemic diarrhea coronavirus (PEDV) causes an extremely high fatality rate in piglets^7^. SARS-CoV-2 emerged from China in 2019^8,9^ and was declared a pandemic during the first quarter of 2020 with an extremely high requirement for a vaccine to be provided in a short timeframe. Spike protein is a multifunctional viral protein found on the outside of the virus particle. It initially binds a host cell receptor though its S1 subunit and fuses viral and host membranes through its S2 subunit. In addition to mediating entry, the spike is a critical determinant of viral host range and a major inducer of host immune responses^10^. Due to the key role of the S protein, it is the main target for antibody-mediated neutralization^11
^.

### 1.2 A brief history of Artificial Intelligence and Deep Learning

Artificial intelligence, based on the assumption that the process of human thought can be mechanized was first conceived as an academic discipline in 1956 following Turing’s landmark paper in 1950^12^. Deep learning (DL) is a subset of Artificial Intelligence that employs deep neural networks requiring the use of a training set and are modelled on the circuitry in the human brain. DL has its roots in Rosenblatt’s Perceptron neural network of 1957-1960^13^. DL algorithms use multiple layers to progressively extract higher level features from raw input. Many different architectures of neural network exist, and for the most part are involved in applications with image recognition. Some recent uses of DL include self-driving cars and robots; with creative projects using DL to compose music, create novel art from different styles of art and write ‘fake news’. The recent advent of much increased processing power and availability in the shape of graphical processing units (GPUs), and CUDA architecture from Nvidia, have paved the way for a renewed interest in machine learning and DL algorithms.

### 1.3 Recurrent Neural Network

The recurrent neural network (RNN) is a type of neural network usually used for text encoding implementations, mainly through whole word encoding and the bag of words concept. Whilst character encoding has been used on occasion this is less often used and is less well supported by software libraries, potentially due to higher memory requirements. The recurrent neural network (RNN)^14^, is trained on a set of sequences using an optimization algorithm with estimations of gradient descent combined with backpropagation through time. The RNN has the potential to consider previously seen data such as the character or word that came before the current time step using units such as long short-term memory cells^15^ (LSTM) or gated recurrent units^16^ (GRU).

### 1.4 Current and upcoming uses for DL in medicine and health care

Recently, uses for DL have been described in health care, particularly in screening for breast cancer^17,18^ and for use with electrocardiogram (ECG) traces^19,20^. Potential novel antibiotics were searched out by screening known drug databases for structures^21^. The strength of this approach lies in that these drugs already have significant results and may have clinical trial data. In 2007, Hochreiter, Heusel and Obermayer proposed the use of LSTM for protein homology detection^15^, commenting that LSTM is capable of automatically extracting local and global sequence statistics like hydrophobicity, polarity, volume and polarizability and combining them with a pattern.

Common bioinformatic techniques involve searching and identifying matches and differences both at small and larger scale and clustering sequences. Hochreiter, Heusel and Obermayer specifically identified the following features during the use of RNNs which would not be otherwise identified using other common bioinformatic techniques.

i. extraction of dependencies between subsequences. A subsequence AB, e.g. may only be indicative if it is followed later by the subsequence CD
ii. extraction of correlations within subsequences. Both AB and CD subsequences may be indicative for the class (motif [AC]– [BD]), however, AD may not be indicative for the class (AD is a negative pattern)
iii. extraction of global sequence characteristics (hydrophobicity or atomic weight) and
iv. extraction of dependencies between amino acids which range over a long interval in the amino acid sequence.

## Methods

### 2.1 Recurrent Neural Network (RNN) Architecture

In creating the model described in this study, character encoding was used on the sequences in the training set. Alternative model architectures were considered and trialled, however, the most successful was composed of a single embedding layer and a gated recurrent unit (GRU) with 1024 RNN units followed by a dense linear layer. The model was trained in Tensorflow 2.1.0^22^ with Keras^23^ using an Adam optimizer^24^ with AMSgrad^25^ option and adaptive learning rate over 15 epochs, where losses fell gradually from an initial 3.259 to 0.266. Python 3.5 and above is required. Figures 1 and 3 were generated in R 3.5.1 (‘Feather Spray’)^26^ with ggplot2^27^ libraries and the Wes Anderson colour palette.

**Figure 1.**
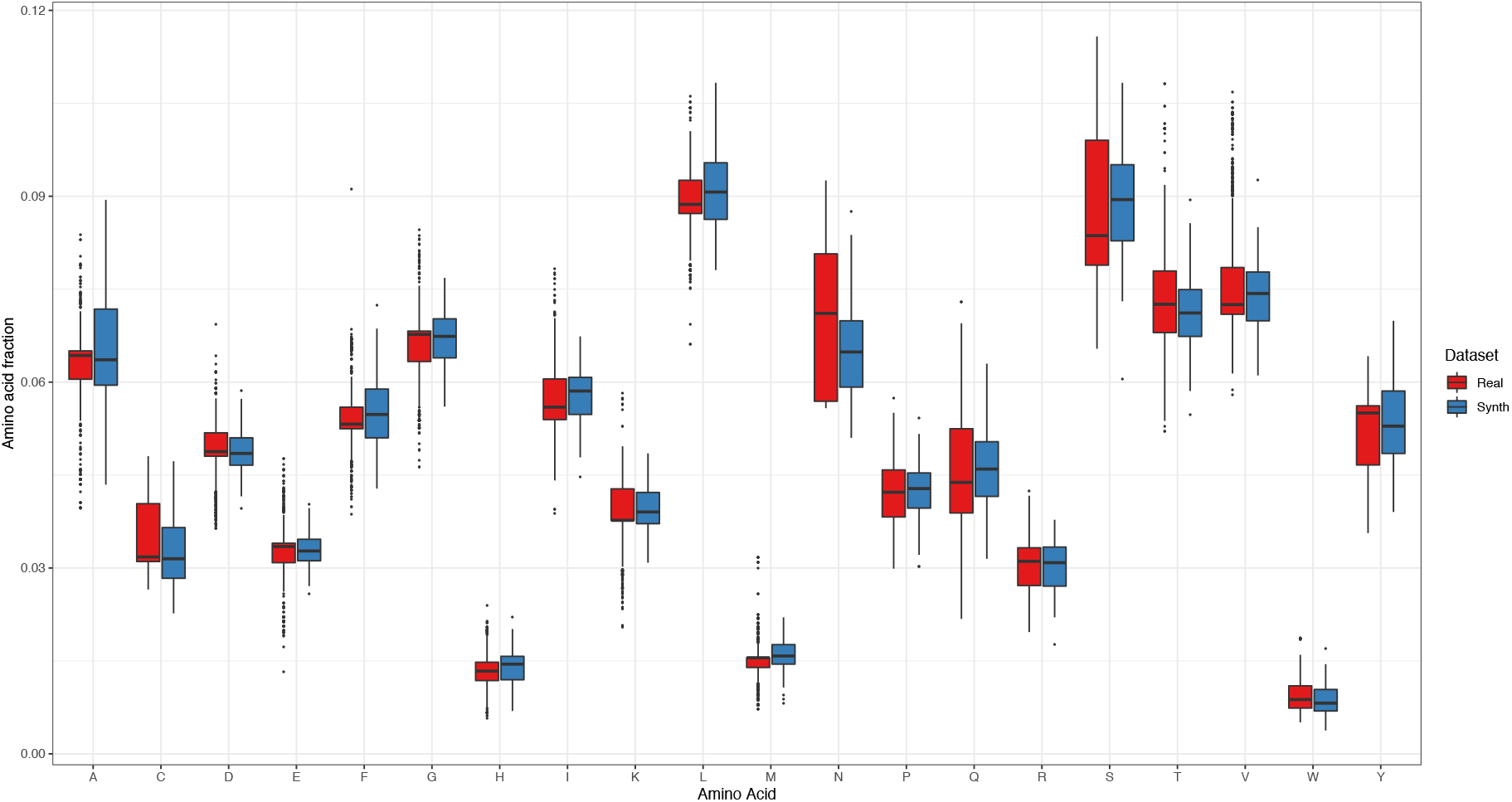
Comparison of the amino acid composition of the real and simulated proteins. Boxplot graph showing the amino acid composition of each amino acid as a fraction of the protein sequence in both the real dataset (red) and the synthesized dataset (blue). The line in the box represents the median. The amino acid single letter code is shown on the X axis with the fraction of the amino acid in each sequence on the y axis as calculated by Biopython ProtParam module. The ‘Real’ dataset comprised 2406 sequences in total whilst the ‘Synth’ simulated example dataset was a sample of 100 sequences.

### 2.2 Coronavirus training set

A training dataset was formulated from a wide variety of coronavirus spike protein sequences from alpha, beta, gamma and delta coronaviruses and constituted isolates from many different animals. The dataset was downloaded from NCBI Genbank on 19^th^ March 2020, prior to any very large release of newly sequenced beta-coronaviruses from SARS-CoV-2, which at the time of writing are destined for later release and stored at Gisaid.org. The total number of spike protein sequences in the final training dataset was 2406, encompassing 511 sequences from Human CoV including examples of SARS-CoV-1 and MERS as well as SARS-CoV-2 (hCoV-19), 232 Bovine, 194 Noctilionine (Bat), 106 Porcine and several samples from other animals including camel, Chinese ferret-badger, hedgehog, dog, deer, avian and whale. Downloaded sequences were searched and cleaned to remove poorer quality and partial sequences and subunits.

## Results

### 3.1 Training dataset manipulation

Newer sequences of SARS-CoV-2 were not included in the training dataset in the belief that the high number of these closely-related beta coronavirus (CoV) sequences could cause bias in the training dataset. Initial processing involved generating a file of 4424729 sequences of 15-mer Kmer windows (overlapping sequences shifted by one amino acid at a time). This is a recognised strategy to help with longer sequences. However, the length of the Kmer was later raised to 100 to improve results and thereafter generated on the fly.

Preliminary investigations on this dataset trained an RNN for hundreds of epochs on GPU with one-hot encoding and with deep LSTM cells. However, a model with a cleaner dataset, a GRU and character indexing proved more successful and relevant in providing simulated sequences. Characteristics of the sequences are examined below.

These downstream sequences were used to investigate the possibility of creating simulated new spike protein sequences that could potentially arise as a chance zoonosis event in future.

### 3.2 Characteristics of DL simulated Spike proteins

To create predictions, the RNN is initially given a short seed protein sequence. The seed sequence can be passed as a random choice from previously sequenced spike proteins or formulated of random choices of amino acids starting with Methionine chosen by the python random library. In this study, the RNN was then able to provide sequences up to the full length of spike protein, a maximum length in the input dataset of 1582 amino acids with a mean length of 1324.4 amino acids. The maximum sequence identity that a simulated sequence achieved in BLAST matches against the training set was 100% sequence identity over 875 amino acids with a temperature scaling value of 1.0 (see below for details) or 100% identity over the full length of the protein with a temperature scaling value of 0.5. The lengths of all the synthesized proteins were fixed at 1588 amino acids. A dataset of 100 DL synthesised spike protein sequences was collected for preliminary investigations. The RNN was initially provided with seed sequences of 16 amino acids chosen at random from the full dataset of spike proteins. The amino acid complement of the real and synthesized spike proteins in the datasets is as compared below in Figure 1. Although the amino acid complements show some differences, it is clear that there are significant similarities across the two datasets. The simulated sequences generated for this figure are provided with the model and source code (as described below).

#### 3.21 Sequence matching

All 100 of the simulated sequences had a significant BLASTP match to Spike protein from one or more coronavirus sequences with BLAST searches of the query sequences against the entire database of NR. A partial example alignment is shown in Figure 2 with a full alignment in Supplementary Data 1.

**Figure 2.**
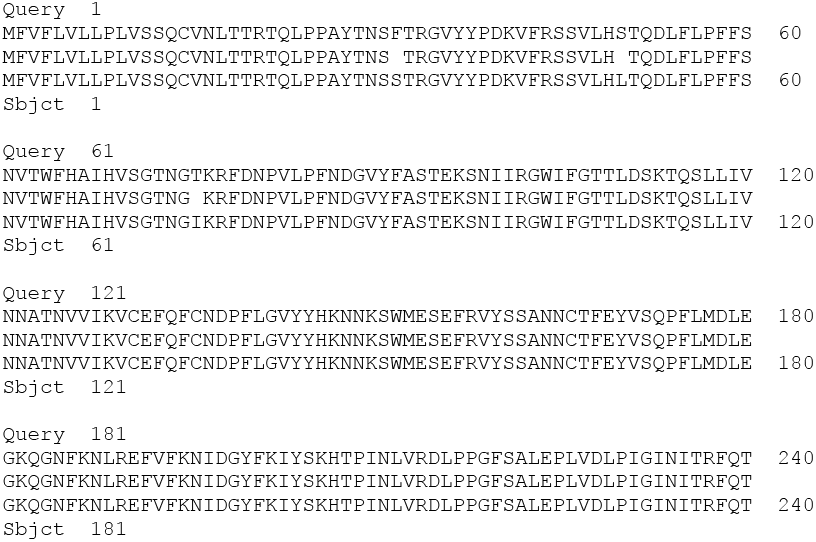
Partial BLAST alignment of Spike protein from a simulated query protein against Bat coronavirus RaTG13^28^. Partial BLASTP alignment of search of a Query (Query) simulated sequence against a member of the training dataset of real spike proteins (Sbjct). In total the identities were 940/1065 (88%) and positives were 1003/1065 (94%). The complete alignment is shown in Supplementary Data 1 (S1).

**Table 1.** Pfam Domain complements in the 100 simulated sequences. Common Pfam domains and their counts identified within the original 100 simulated sequences which are also found in real Spike proteins, showing that all 100 sequences had C-terminal Spike domains. Other domains were identified in full Pfam-A but the most common were Corona_S2, Spike_rec_bind and Spike_NTD. Database Pfam-A.SARS-CoV-2 refers to the April 2, 2020 update for SARS-CoV Pfam domains (Xfam Blog https://xfam.wordpress.com/2020/04/02/pfam-sars-cov-2-special-update/)

| Pfam domain | Pfam DB | Full Name | Count |
| --- | --- | --- | --- |
| Corona_S2 | Pfam-A | Coronavirus S2 glycoprotein | 100 |
| Corona_S1 | Pfam-A | Coronavirus S1 glycoprotein | 13 |
| Spike_rec_bind | Pfam-A | Spike receptor binding domain | 81 |
| Spike_NTD | Pfam-A | Spike glycoprotein N-terminal | 53 |
| CoV_NSP2_C | Pfam-A.SARS-CoV-2 | Coronavirus replicase NSP2, C-terminus | 6 |
| CoV_S1_C | Pfam-A.SARS-CoV-2 | Coronavirus Spike S1, C-terminus | 75 |
| bCoV_S1_RBD | Pfam-A.SARS-CoV-2 | Betacoronavirus Spike S1, receptor-binding | 82 |
| bCoV_S1_N | Pfam-A.SARS-CoV-2 | Betacoronavirus-like spike S1, N-terminus | 88 |
| CoV_S2 | Pfam-A.SARS-CoV-2 | Coronavirus Spike glycoprotein S2 | 100 |

**Figure 3.**
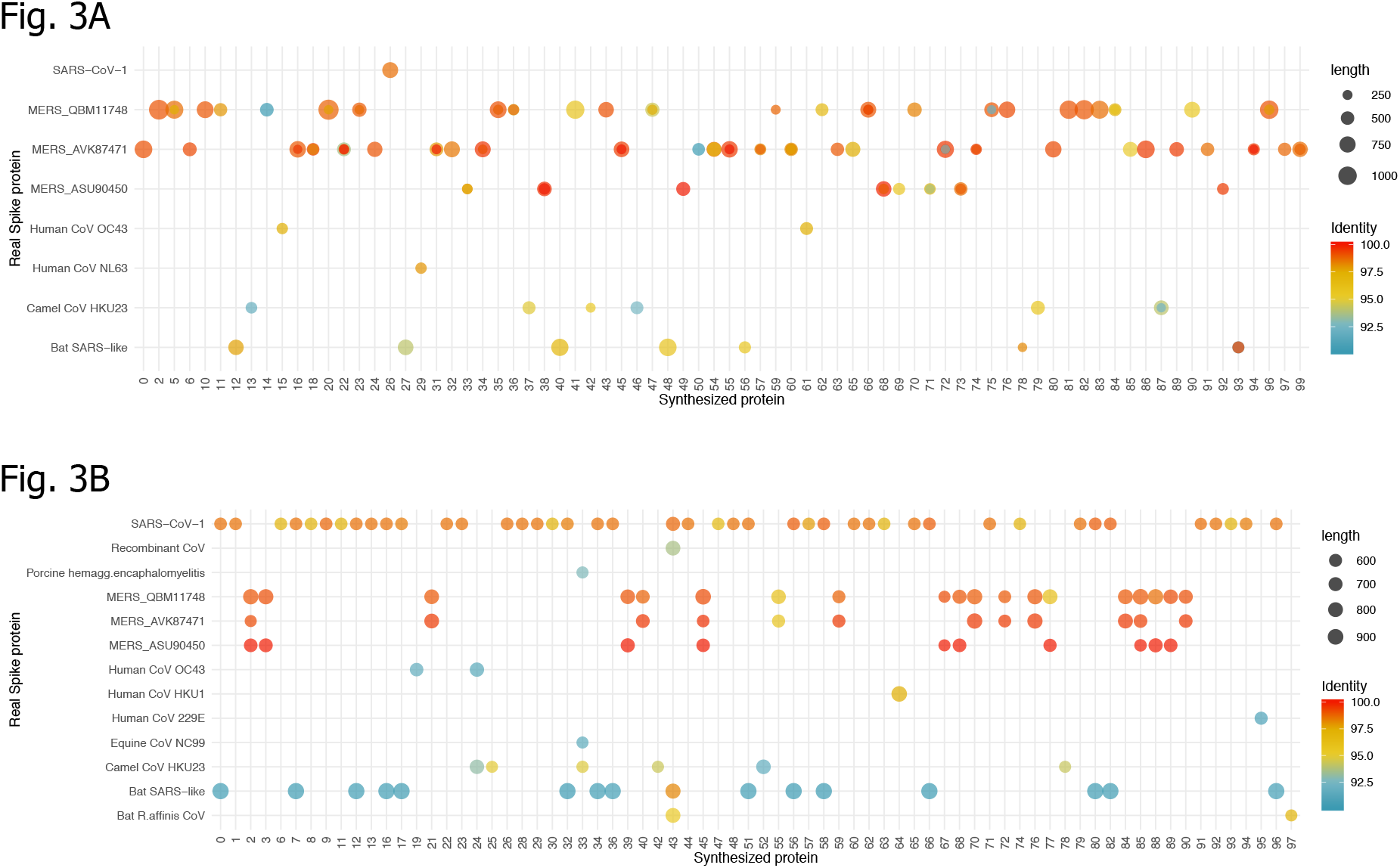
BLAST matches of simulated sequences against known Spike proteins. **BLAST searches of DL synthesized sequences against a clustered version of the Spike protein sequence training set.** 3A shows BLAST searches of the original 100 DL synthesized sequences filtered for matches by length over 200bp and identity over 90%. 3B shows the second set of DL synthesized sequences which were all given identical feeder sequence of 64 amino acids from the start of SARS-CoV-2. This graph is filtered for matches by length over 500 bp and identity over 90%. Highest length matches are represented by the largest diameter circle and most red colour. However, large circles are generally of interest since the identity cut-off is high. Unfiltered data is presented in Supplementary Figure 1.

The real spike protein training dataset was clustered and deduplicated using cd-hit with default parameters resulting in 154 clusters. The 100 simulated sequences were searched with BLASTP against the representative cluster sequences. Figure 3A shows that the best BLAST hits for the first set of simulated sequences covered several distinct clusters. There were several hits to MERS clusters, possibly due to a high representation of MERS in the training set. Figure 3B shows the equivalent BLAST hits on the second set of 100 simulated sequences that each had an identical seed text of 64 amino acids from the start of SARS-CoV-2 spike. There are a high number of SARS-CoV matches, as well as Bat SARS-like sequences, although some samples still shared high identities with MERS sequences.

#### 3.22 Pfam domain complements of simulated protein sequences

Significant Pfam domain hits were uncovered on searching the query sequences with HMMER3^29^ using hmmsearch against the Pfam-A^30^ database. HMMER3 searches of the synthesized proteins against Pfam_A.hmm database revealed Pfam domains that were expected within a coronavirus spike protein (below).

This compared favourably with the most commonly found domains within the real training dataset of 2504 proteins which had 1781 domains of Spike_rec_bind, 2413 Corona_S2, 1052 Spike_NTD and 502 Corona_S1 (Coronavirus S1 glycoprotein domain) among others. According to Pfam architectures^30^, domain Corona_S2 is found in real spike proteins in the databases with either Corona_S1, Spike_NTD and Spike_rec_bind, just Spike_rec_bind, with Spike_NTD and 2 × Spike_rec_bind and in some sequences as a standalone domain.

#### 3.33 Prediction temperature parameter

During prediction, probabilities are generated for the next character in the sequence of the amino acid single letter alphabet. A parameter known as the ‘temperature’ of 0.5 produces more similar sequences by scaling the resultant probabilities of the multinomial distribution, for example the model was able to reach 100% identity over the full length of Spike SARS glycoprotein at a temperature of 0.5 with the only differences being in the seed text. The purpose of this study is to provide sequences that are not identical to known sequences hence we may find better use of a higher temperature value. A second dataset sample of 100 synthesized sequences was formed by specifically using a seed text of 64 amino acids from the SARS-CoV-2 spike protein for each simulated sequence. When simulated sequences were clustered with cd-hit^31^ at the default 90% level of identity, the dataset provided 51 separate clusters in which Cluster 0 had 27 members ranging from 92% - 100% identity which corresponded to SARS-CoV-1 type, Cluster 1 had 13 members of 97-100% identity which corresponded to Bat RaTG13/SARS-CoV-2 type, Cluster 38 had 3 members corresponding to MERS type, Clusters 2, 8 and 8 each had two members, each example of the rest of the dataset clustered separately. Therefore, the seed text provided SARS-like hits in several but not all cases.

Once the initial predicted protein is finished, the prediction commences a new protein again immediately if the maximum number of characters has not been reached. Therefore, in some cases there were hybrid matches to parts of sequence from spike proteins in the dataset.

Some sequences were definitively of interest to this study, such as a synthesized protein with 97% full length identity to a Bat beta-coronavirus sequence isolated from *Chaerephon plicata* in Yunnan in 2011^32^. Further sequences of interest included those with high identity over stretches of the protein sequence to SARS-CoV-1 or SARS-CoV-2, particularly those including hybrid regions. This dataset is provided with the model and source (as described below).

With the study of the most relevant sequences of 2020, SARS-CoV-1 and SARS-CoV-2 in mind, a much larger dataset of 1000 simulated sequences was generated with the same SARS-CoV seed text as previously. These simulated sequences were clustered with cd-hit and clusters corresponding to SARS-CoV-1 and SARS-CoV-2 were aligned together with examples of the real spike proteins from SARS-CoV-1 and SARS-CoV-2. Alignments were sliced in Biopython to remove extra sequences at the ends and the seed text at the start. The resulting multiple sequence alignment is shown in Supplementary Figure 2. This alignment indicates that residues important in human ACE2 recognition^33^ are broadly conserved across the simulated sequences.

## Conclusions

This study used a comprehensive training set formulated from Coronavirus Spike protein sequences in the sequence databases for DL neural networks to produce novel sequence from a short feeder seed text. The novel sequences share features that can be searched with bioinformatics tools to bring out highly significant BLAST matches and Pfam domain matches. That each of the sequences examined had BLAST and Pfam matches to Spike protein is exciting and warrants further consideration.

Interestingly, in an example of a very accurate alignment, the prediction query was able to fill in a blank amino acid (G) where one was called as X in the real sequence that exactly matched other sequences of that type. It is to be noted that the generation of very large numbers of these synthesized sequences from the trained data is trivial and many thousands can be further generated and examined from the model (model & source code available at: https://github.com/LCrossman). It is hoped that the simulated sequences may have the potential to aid in forthcoming vaccine searches effectively by allowing a peek into a fast-forward of evolution to see what sequences may possibly arise in future. However, it is to be noted that the training set is only as comprehensive as the initial database, animal CoV sequences may exist elsewhere that are not present in our database. Additionally, biases may exist, for example, the training set is overrepresented in MERS sequences and these are also heavily represented in the random simulated sequences. In some cases the resultant protein may not represent a viable protein (there could be stability or other issues). One additional potential issue could be that the premise is difficult to test.

Nonetheless, the potential production of novel sequence by DL is exciting and warrants further consideration.

## Acknowledgements

The author thanks Xfam for making available the Pfam-A SARS update *via* their blog and all key workers of the COVID-19 pandemic.

