## Supplementary Data 1_S1 for "Leveraging Deep Learning to Simulate Coronavirus Spike proteins has the potential to predict future Zoonotic sequences"

BLAST alignment for example query simulated sequence best hit against NR database

Spike glycoprotein [Bat coronavirus RaTG13]

Sequence ID: QHR63300.2Length: 1269Number of Matches: 1

Range 1: 1 to 1065GenPeptGraphicsNext MatchPrevious Match

| Score | Expect | Method | Identities | Positives |
|---|---|---|---|---|
| 1966 bits(5093) | 0.0 | Compositional matrix adjust. | 940/1065(88%) | 1003/1065(94%) |

```
Query  1    MFVFLVLLPLVSSQCVNLTTRTQLPPAYTNSFTRGVYYPDKVFRSSVLHSTQDLFLPFFS  60
            MFVFLVLLPLVSSQCVNLTTRTQLPPAYTNS TRGVYYPDKVFRSSVLH TQDLFLPFFS
Sbjct  1    MFVFLVLLPLVSSQCVNLTTRTQLPPAYTNSSTRGVYYPDKVFRSSVLHLTQDLFLPFFS  60

Query  61   NVTWFHAIHVSGTNGTKRFDNPVLPFNDGVYFASTEKSNIIRGWIFGTTLDSKTQSLLIV  120
            NVTWFHAIHVSGTNG KRFDNPVLPFNDGVYFASTEKSNIIRGWIFGTTLDSKTQSLLIV
Sbjct  61   NVTWFHAIHVSGTNGIKRFDNPVLPFNDGVYFASTEKSNIIRGWIFGTTLDSKTQSLLIV  120

Query  121  NNATNVVIKVCEFQFCNDPFLGVYYHKNNKSWMESEFRVYSSANNCTFEYVSQPFLMDLE  180
            NNATNVVIKVCEFQFCNDPFLGVYYHKNNKSWMESEFRVYSSANNCTFEYVSQPFLMDLE
Sbjct  121  NNATNVVIKVCEFQFCNDPFLGVYYHKNNKSWMESEFRVYSSANNCTFEYVSQPFLMDLE  180

Query  181  GKQGNFKNLREFVFKNIDGYFKIYSKHTPINLVRDLPPGFSALEPLVDLPIGINITRFQT  240
            GKQGNFKNLREFVFKNIDGYFKIYSKHTPINLVRDLPPGFSALEPLVDLPIGINITRFQT
Sbjct  181  GKQGNFKNLREFVFKNIDGYFKIYSKHTPINLVRDLPPGFSALEPLVDLPIGINITRFQT  240

Query  241  LLALHRSYLTPGDSSSGWTAGAAAYYVGYLQPRTFLLKYNENGTITDAVDCALDPLSETK  300
            LLALHRSYLTPGDSSSGWTAGAAAYYVGYLQPRTFLLKYNENGTITDAVDCALDPLSETK
Sbjct  241  LLALHRSYLTPGDSSSGWTAGAAAYYVGYLQPRTFLLKYNENGTITDAVDCALDPLSETK  300

Query  301  CTLKSFTVEKGIYQTSNFRVQPTESIVRFPNITNLCPFGEVFNATTFPSVYAWERKRISN  360
            CTLKSFTVEKGIYQTSNFRVQPT+SIVRFPNITNLCPFGEVFNATTF SVYAW RKRISN
Sbjct  301  CTLKSFTVEKGIYQTSNFRVQPTDSIVRFPNITNLCPFGEVFNATTFASVYAWNRKRISN  360

Query  361  CVADYSVLYNSTSFSTFKCYGVSATKLNDLCFSNVYADSFVVKGDDVRQIAPGQTGVIAD  420
            CVADYSVLYNSTSFSTFKCYGVS TKLNDLCF+NVYADSFV+ GD+VRQIAPGQTG IAD
Sbjct  361  CVADYSVLYNSTSFSTFKCYGVSPTKLNDLCFTNVYADSFVITGDEVRQIAPGQTGKIAD  420

Query  421  YNYKLPDDFMGCVLAWNTRNIDATSTGNHNYKYRYLRHGKLRPFERDISNVPFSPDGKPC  480
            YNYKLPDDF GCV+AWN+++IDA   GN NY YR  R   L+PFERDIS   +    KPC
Sbjct  421  YNYKLPDDFTGCVIAWNSKHIDAKEGGNFNYLYRLFRKANLKPFERDISTEIYQAGSKPC  480

Query  481  T-PPALNCYWPLNDYGFYTTTGIGYQPYRVVVLSFELLNAPATVCGPKLSTDLIKNQCVN  539
                 LNCY+PL  YGFY T G+G+QPYRVVVLSFELLNAPATVCGPK ST+L+KN+CVN
Sbjct  481  NGQTGLNCYYPLYRYGFYPTDGVGHQPYRVVVLSFELLNAPATVCGPKKSTNLVKNKCVN  540
```

```
Query  540   FNFNGLTGTGVLTPSSKRFQPFQQFGRDVSDFTDSVRDPKTSEILDISPCSFGGVSVITP  599
             FNFNGLTGTGVLT S+K+F PFQQFGRD++D TD+VRDP+T EILDI+PCSFGGVSVITP
Sbjct  541   FNFNGLTGTGVLTESNKKFLPFQQFGRDIADTTDAVRDPQTLEILDITPCSFGGVSVITP  600

Query  600   GTNASSEVAVLYQDVNCTDVSTAIHADQLTPAWRIYSTGNNVFQTQAGCLIGAEHVDTSY  659
             GTNAS++VAVLYQDVNCT+V  AIHADQLTP WR+YSTG+NVFQT+AGCLIGAEHV+ SY
Sbjct  601   GTNASNQVAVLYQDVNCTEVPVAIHADQLTPTWRVYSTGSNVFQTRAGCLIGAEHVNNSY  660

Query  660   ECDIPIGAGICASYHTVSLLRSTSQKSIVAYTMSLGADSSIAYSNNTIAIPTNFSISITT  719
             ECDIPIGAGICASY T +  RS + +SI+AYTMSLGA++S+AYSNN+IAIPTNF+IS+TT
Sbjct  661   ECDIPIGAGICASYQTQTNSRSVASQSIIAYTMSLGAENSVAYSNNSIAIPTNFTISVTT  720

Query  720   EVMPVSMAKTSVDCNMYICGDSTECANLLLQYGSFCTQLNRALSGIAAEQDRNTREVFAQ  779
             E++PVSM KTSVDC MYICGDSTEC+NLLLQYGSFCTQLNRAL+GIA EQD+NT+EVFAQ
Sbjct  721   EILPVSMTKTSVDCTMYICGDSTECSNLLLQYGSFCTQLNRALTGIAVEQDKNTQEVFAQ  780

Query  780   VKQMYKTPTLKYFGGFNFSQILPDPLKPTKRSFIEDLLFNKVTLADAGFMKQYGECLGDI  839
             VKQ+YKTP +K FGGFNFSQILPDP KP+KRSFIEDLLFNKVTLADAGF+KQYG+CLGDI
Sbjct  781   VKQIYKTPPIKDFGGFNFSQILPDPSKPSKRSFIEDLLFNKVTLADAGFIKQYGDCLGDI  840

Query  840   NARDLICAQKFNGLTVLPPLLTDDMIAAYTAALVSGTATAGWTFGAGAALQIPFAMQMAY  899
              ARDLICAQKFNGLTVLPPLLTD+MIA YT+AL++GT T+GWTFGAGAALQIPFAMQMAY
Sbjct  841   AARDLICAQKFNGLTVLPPLLTDEMIAQYTSALLAGTITSGWTFGAGAALQIPFAMQMAY  900

Query  900   RFNGIGVTQNVLYENQKQIANQFNKAISQIQESLTTTSTALGKLQDVVNQNAQALNTLVK  959
             RFNGIGVTQNVLYENQK IANQFN AI +IQ+SL++T++ALGKLQDVVNQNAQALNTLVK
Sbjct  901   RFNGIGVTQNVLYENQKLIANQFNSAIGKIQDSLSSTASALGKLQDVVNQNAQALNTLVK  960

Query  960   QLSSNFGAISSVLNDILSRLDKVEAEVQIDRLITGRLQSLQTYVTQQLIRAAEIRASANL  1019
             QLSSNFGAISSVLNDILSRLDKVEAEVQIDRLITGRLQSLQTYVTQQLIRAAEIRASANL
Sbjct  961   QLSSNFGAISSVLNDILSRLDKVEAEVQIDRLITGRLQSLQTYVTQQLIRAAEIRASANL  1020

Query  1020  AATKMSECVLGQSKRVDFCGKGYHLMSFPQAAPHGVVFLHVTYVP  1064
             AATKMSECVLGQSKRVDFCGKGYHLMSFPQ+APHGVVFLHVTYVP
Sbjct  1021  AATKMSECVLGQSKRVDFCGKGYHLMSFPQSAPHGVVFLHVTYVP  1065
```
