## Supplementary Figure 1_SF1 for "Leveraging Deep Learning to Simulate Coronavirus Spike proteins has the potential to predict future Zoonotic sequences"

### SF1A  BLASTP matches for the initial set of simulated query proteins against a clustered version of the Spike protein training set

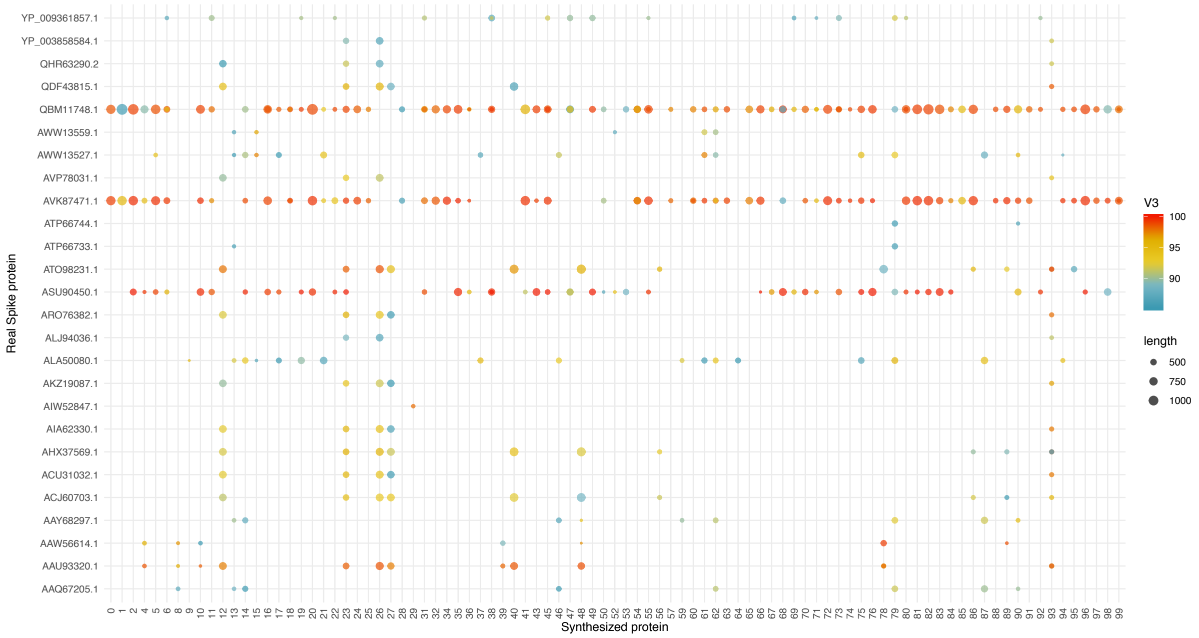


The model RNN was given a seed text of 16 amino acids randomly selected from the full training set of Spike proteins.  BLAST matches were filtered to remove those of length less than 500 bp and sequence identity of less than 80%.

### SF1B  BLAST matches for second set of query proteins against the clustered training set Spike proteins

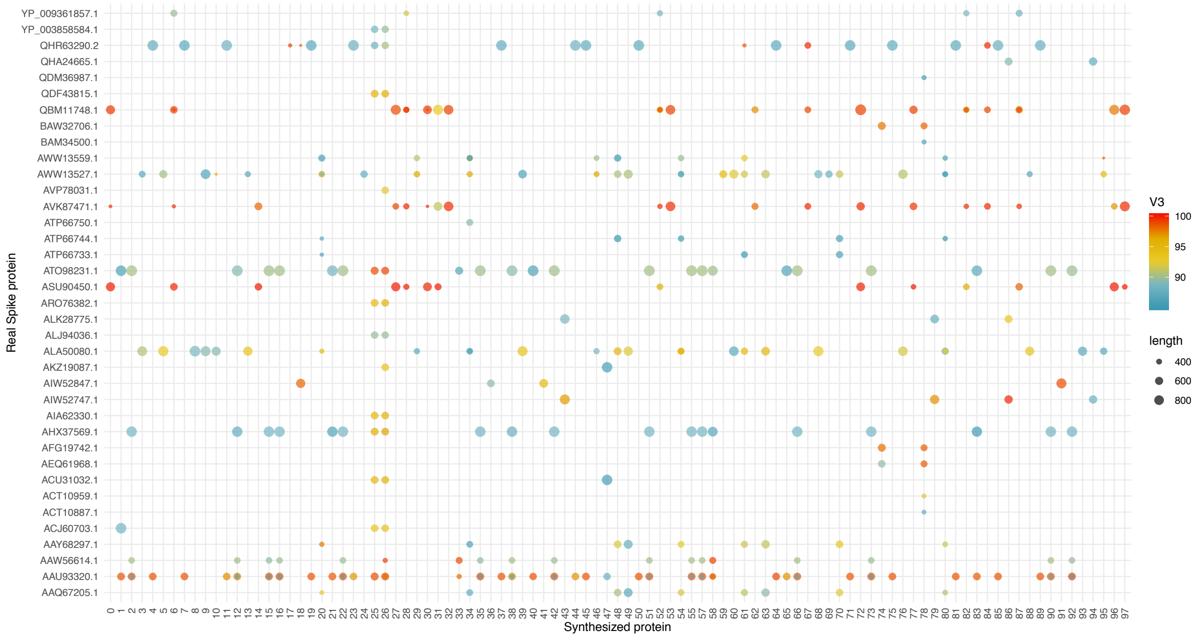


BLAST matches were filtered to remove any of less than 400 bp and of less than 80% identity.  The seed text was an identical sequence of 64 amino acids from SARS-CoV-2 in each case:
MFVFLVLLPLVSSQCVNLTTRTQLPPAYTNSFTRGVYYPDKVFRSSVLHSTQDLFLPFFSNVTWF
