## Supplementary figures and images for "Leveraging Deep Learning to Simulate Coronavirus Spike proteins has the potential to predict future Zoonotic sequences"

### Supplementary Figure 1_SF2

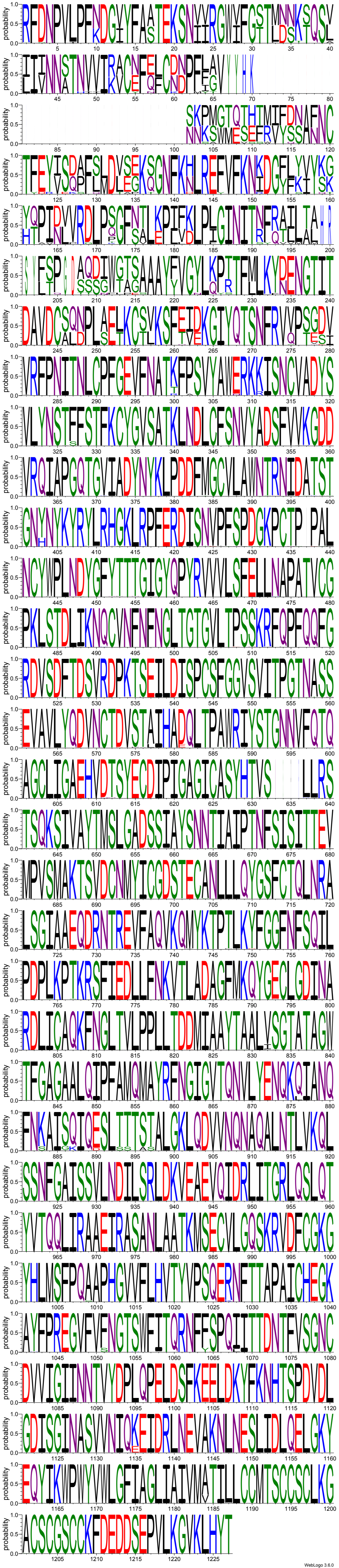
